# Reactivating ordinal position information from auditory sequence memory in human brains

**DOI:** 10.1101/2022.04.04.487064

**Authors:** Ying Fan, Huan Luo

**Affiliations:** School of Psychological and Cognitive Sciences, Peking University; PKU-IDG/McGovern Institute for Brain Research, Peking University; Beijing Key Laboratory of Behavior and Mental Health, Peking University

**Keywords:** auditory working memory, sequence memory, ordinal position, reactivation, impulse-response, EEG

## Abstract

Retaining a sequence of events in their order is a core ability of many cognitive functions, such as speech recognition, movement control, and episodic memory. Although content representations have been widely studied in working memory (WM), little is known about how ordinal position information of an auditory sequence is retained in the human brain as well as its coding characteristics. In fact, there is still a lack of an efficient approach to directly access the stored ordinal position code, since the neural activities enter a relatively ‘activity-silent’ WM state during WM retention. Here participants performed an auditory sequence WM task with their brain activities recorded using electroencephalography (EEG). We developed new triggering events that could successfully reactivate neural representations of ordinal position from the ‘activity-silent’ retention stage. Importantly, the rank reactivation is further related to recognition behavior, confirming its indexing of WM storage. Furthermore, the ordinal position code displays an intriguing ‘stable-dynamic’ format, i.e., undergoing the same dynamic neutral trajectory during both encoding and retention (whenever reactivated). Overall, our results provide an effective approach to access the behaviorally-relevant ordinal position information in auditory sequence WM and reveal its new temporal characteristics.

## Introduction

How does the human brain store auditory sequence, the most typical auditory experience (e.g., sentence, melody, etc.) in our daily lives? This process relies on the correct ordering of auditory contents in working memory (WM). In other words, the ordinal position for each item in the sequence needs to be well represented and stored, without which our memory would be just fragmented, unorganized pieces. Amounts of previous studies have focused on the neural representation of content information (e.g., tone frequency, auditory category, spatial location, etc.) in WM (Albouy et al., 2017; Bays et al., 2009; Harrison & Tong, 2009; Jonides et al., 1993), yet the maintenance mechanism for ordinal position information remains obscure.

How to access information retained in WM, especially during the delay period? This issue poses a tremendous challenge to human studies using non-invasive recordings since the to-be-memorized information couldn’t be easily or reliably accessed during retention (Lundqvist et al., 2018; Sreenivasan et al., 2014), somewhat commensurate with the ‘activity-silent’ WM view (Stokes, 2015). Recently, an impulse-response approach (Fan et al., 2021; Wolff et al., 2017) has been developed to successfully reactivate WM information during the delay period, thereby the underlying neural mechanism could be studied. For instance, our previous work (Fan et al, 2021) showed that when participants retained a list of pure tones, an auditory white noise that was neither memory-related nor task-relevant could trigger neural representations of memorized tone pitch, which further predicts WM behavior. The white-noise impulse is posited to act as a triggering event to transiently perturb the WM system so that the retained information could be reactivated (Manohar et al., 2019). Meanwhile, this was not the case for the ordinal position, which failed to be activated by the neutral white-noise impulse. This implies that the pitch and ordinal position of auditory sequences are retained in different ways and would thus be triggered by different neutral impulses. Therefore, it is crucial to develop triggering events, akin to the white-noise auditory impulse for pitch, to probe the storage mechanism of ordinal position in auditory WM.

Another puzzle is about the coding characteristics of ordinal position in WM. Prior studies demonstrate that content representation, such as orientation, tone pitch, etc., undergoes a dynamic transformation from encoding to maintenance (Kamiński & Rutishauser, 2020; Rademaker et al., 2019; Yu et al., 2020). In contrast, the structure information tends to be coded in a stable format (Bernardi et al., 2020; Kalm & Norris, 2017; Luyckx et al., 2019; Walsh, 2003). Meanwhile, there still lacks clear time-resolved evidence characterizing its representational dynamics. For example, the structure representation might be a completely stationary format and keep the same over time (“stable-stable”). Alternatively, it might follow a dynamic trajectory in neural space but repeat the same trajectory when reactivated (“stable-dynamic”). The latter alternative would also predict encoding-to-maintaining representational generalization for structure, as demonstrated in previous studies (Fan et al., 2021; Kalm et al., 2017). Unraveling the coding dynamics is also critical for our understanding of ordinal position representation in WM.

Here in an auditory sequence WM experiment, human participants were asked to retain a 3-tone auditory sequence and tested on the ordinal position of one of the three memory tones. Their brain activities were recorded using electroencephalogram (EEG) and a multivariate decoding approach was employed to decode the ordinal position information. Critically, we designed a new type of neutral impulse (i.e., a visual symbolic image) which could reactivate the ordinal position of memory tones during retention and the neural reactivations are further related to memory performance. Moreover, the ordinal position code displays an interesting “stable-dynamic” representational format throughout the WM process, i.e., following a neutral trajectory over time and repeating the same dynamic trajectory whenever reactivated. Overall, our study provides an efficient approach to probe behaviorally-relevant rank information retained in auditory sequence WM and demonstrate its stable-dynamic characteristics.

## Methods

### Participants

Thirty-one (15 females and 16 males, mean age 21.8, range 18-26 years) healthy participants were recruited in the current research, after providing written informed consent. All participants had normal or corrected-to-normal vision, with no history of psychiatric or neurological disorders. Participants received compensation for participation. The experiment was approved by the Departmental Ethical Committee of Peking University.

### Apparatus and Stimuli

The whole experiment was controlled by Psychtoolbox under MATLAB. Visual stimuli were presented on a Display ++ LCD screen with a resolution of 1920 by 1080 pixels running at a refresh rate of 120 Hz. Auditory stimuli, whose intensities were approximately 65 dB (62.1 to 67.3 dB) SPL, were pre-generated by customized MATLAB codes and played via a Sennheiser CX300S earphone through an RME Babyface pro external sound card during the experiment. Participant’s head was stabilized by a chin rest, which was fixed at 60 cm away from the screen. Responses were collected using a USB keyboard.

### Experimental procedure

The presentation of a cross (0.9° visual angle) in the center of the screen indicated the start of a trial. This cross stayed in the center throughout the whole trial except during the neutral impulse presentation and the ordinal position report screen (Figure 1A left). Participants were instructed to fixate at this central cross and keep the eye blink to the minimum within each trial. During the encoding period, three memory tones (pure tones) with different frequencies that were pseudo-randomly selected from a uniform distribution of 5 frequencies (420 Hz, 593 Hz, 840 Hz, 1187 Hz, and 1689 Hz) were presented sequentially. The duration of each memory tone was 150 ms including 10 ms ramp up and 10 ms ramp down time. Participants were required to memorize these three tones’ frequencies in their presented order. After a delay of 2000 ms, a retro-cuing tone, which had the same frequency as one of the memory tones was presented. Participants maintained the ordinal position of the retro-cuing tone until response period. Crucially, during the delay period, a 100-ms visual neutral impulse – a white circle (visual angle: 18°) embedded with a symbolic ‘0’ – was presented 2100 ms after the retro-cuing tone, aiming to perturb the WM network so that the stored ordinal position might be reactivated. Finally, after another 800 ms interval, the response screen was presented and participants reported the ordinal position. To avoid early motor response preparation after the retro-cuing tone, the correspondence between ordinal positions and response keys was randomly set for each trial. Next trial started 1200 - 1500 ms after the response. There were 540 trials in total, and participants entered a forced 1-2 minutes break session every 30 trials. The whole experiment lasted around 3 hours for each subject.

**Figure 1.**
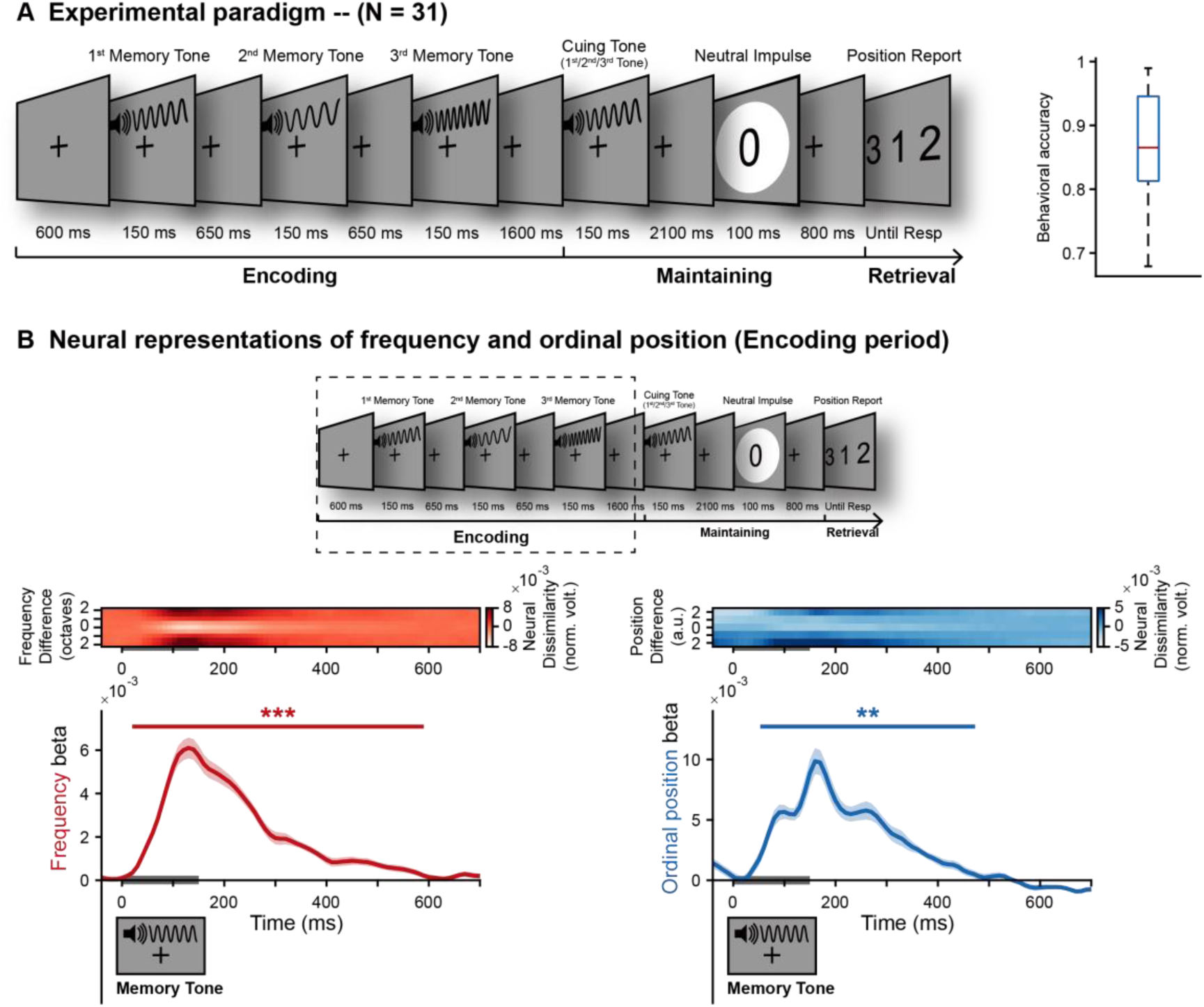
Experimental paradigm and neural representations of pitch and ordinal position during encoding. (A) Experimental paradigm and behavioral results. Left: in each trial, participants needed to memorize a 3-tone sequence (3 frequencies as well as their ordinal positions in the sequence) which consists of three serially presented tones with different frequencies (selected from 420 Hz, 593 Hz, 840 Hz, 1187 Hz, and 1689 Hz). During the maintaining period, a cuing tone having the same frequency as one of the three tones was presented and participants needed to retrieve and hold its corresponding ordinal position (1^st^, 2^nd^, 3^rd^) and make response during subsequent recalling period. Crucially, a visual neutral impulse, i.e., a white circle containing a symbolic ‘0’, was shown on the screen following the cuing tone, aiming to transiently perturb WM system so that memory information could be potentially reactivated. Right: Mean accuracy of ordinal position report (86.88% ±1.37%). (B) Neural representations of tone frequency and ordinal position during encoding period. Upper: dotted box indicates the encoding period for decoding analysis. The responses for the three tones are extracted and analyzed together. Middle: grand average (N = 31) time-resolved neural representational dissimilarity (mean-centered) as a function of physical dissimilarity (y-axis) over time (x-axis) after onset of each tone, for auditory frequency (left) and ordinal position (right). Lower: grand average (N = 31, mean ± SEM) time-resolved decoding performance (regression between neural representational dissimilarity and physical dissimilarity) for tone frequency (left) and ordinal position (right), after onset of each tone. Gray horizontal bar indicates tone presentation. (Shaded area represents ± 1 SEM across participants. ***: p < 0.001; **: p < 0.05; corrected using cluster-based permutation test, cluster-forming threshold p < 0.05).

### EEG acquisition and pre-processing

The EEG signals were acquired using a 64-channel EasyCap. Data was recorded at 500 Hz using two BrainAmp amplifiers (Brain Products) and BrainVision Recorder software (Brain Products). The impedance of all electrodes was kept below 10 kΩ. Offline pre-processing was conducted using fieldtrip (Oostenveld et al., 2011). The EEG data were first re-referenced to the average value of all channels, down-sampled to 100 Hz, and bandpass filtered with the range from 1 Hz to 30 Hz. Independent component analysis was performed to remove eye movement and other artifactual components and the remaining components were then back-projected to the EEG channel space. Data were epoched 500 ms before the first memory tone’s onset to 700 ms after the report screen’s offset. Lastly, epochs with extremely high noises by visual inspection were manually excluded.

### Time-resolved multivariate decoding - RSA analysis

We conducted an 8-fold cross-validated multivariate decoding analysis. First, to remove possible interference of slow trend on decoding results, the mean activities over all trials for each channel were calculated and smoothed by a 150-ms window, which was then subtracted from the original data in each trial (Grootswagers et al., 2017). Moreover, the variance that could be explained by the global field power (GFP) was removed from the original signals for the encoding period data using a cross-validated confound regression method (Snoek et al., 2019).

Both spatial and temporal information were included for the multivariate decoding analysis. Specifically, at each time point, the 64-channel EEG signals at the current as well as previous four time points were included as features (i.e., 64*5 = 320 features). We used an 8-fold cross-validated approach to conduct the decoding analysis by dividing all trials into 8 folds and using one of them as testing dataset and the other 7 folds as training dataset iteratively. We first estimated the condition-specific spatiotemporal activities (i.e., 320 features) by averaging trials belonging to the same condition for the current factor-of-interest using training dataset. Specifically, if the current factor-of-interest is ordinal position, there exist 3 conditions (i.e., 1^st^, 2^nd^, and 3^rd^) and if it is frequency, there are 5 conditions (i.e., 420 Hz, 593 Hz, 840 Hz, 1187 Hz, and 1689 Hz). Next, we calculated the representational dissimilarity (Mahalanabis distance) in the neural spatiotemporal activities between each testing trial and the averaged condition-specific activities at each time point for each factor-of-interest separately. In the meantime, the physical dissimilarity between each testing trial and training conditions was also calculated. For each of the 8 iterations, the neural representational dissimilarity was regressed using the physical dissimilarity. The correlation strength (β) was averaged across all testing trials, yielding the decoding accuracy time course, for pitch and ordinal position, separately, in each participant.

To avoid possible partition bias, the above mentioned 8-fold cross-validation process was repeated 50 times with random partitions. The trial number for each factor-of-interest condition across all participants was set to be equal, chosen as the minimum number of trials of all factor-of-interest conditions across participants. The same trial numbers also guarantee the fair comparisons between high- and low-performance groups (details in latter). A Gaussian-weighted (window length = 40 ms) smooth was further applied on the final decoding time-course.

A non-parametric sign-permutation test (Maris & Oostenveld, 2007) was used to perform statistical tests on the decoding time courses. Specifically, the sign of the correlation strength (β) for each participant at each time point was flipped 100,000 times randomly, through which the null distribution of population mean β was obtained, from which we could estimate the p-value of the observed β. To correct the multiple comparisons over time, we performed a cluster-based permutation test (cluster-forming threshold p < 0.05) and seek the corrected temporal clusters (p < 0.05 or p < 0.001).

### Behavioral relevance

We separated participants into high-performance (top 50%) and low-performance (bottom 50%) groups based on their behavioral accuracies and computed their respective decoding results. After obtaining the time course of decoding results for each performance group, we summed all the beta values reaching significant statistical levels for each group, and conducted a Mann-Whitney U test for comparisons between groups.

### Cross-time generalization analysis and permutation tests

We performed the same RSA-based multivariate decoding analysis as before, except that the representational similarity was calculated on neural activities occurring at different time points. Smoothing with a 2-D Gaussian smoothing kernel with standard deviation of 2 was further applied on the generalization results.

To examine which patterns (Figure 4A), i.e., “stable-sustained”, “stable-dynamic”, or “transient”, could best characterize the generalization results, we developed a new testing method. For “stable-sustained” coding, the cross-time generalization result would show a horizontally-oriented rectangle in shape (Figure 4A left), i.e., stable over time. In contrast, the “stable-dynamic” coding would display a generalization pattern parallel to the diagonal, which could be depicted by a diagonally-oriented parallelogram (Figure 4A middle). To access which coding format could capture the observed results, we first established all possible horizontally-oriented rectangles (testing “stable-sustained”) and diagonally-oriented parallelograms (testing “stable-dynamic) to cover the whole generalization matrix (0-700 ms, 71*71). Furthermore, the center of these rectangles or parallelograms was confined in the lower triangular of the generalization matrix to reduce computational load. As a result, 1677060 horizontally-oriented rectangles and 1677060 diagonally-oriented parallelograms were formed and tested. Next, the beta values enclosed by each of these quadrilaterals were summed to represent their respective characterization performance (Figure 4BCD, middle).

A permutation test was then conducted to test the statistical significance. Specifically, the raw generalization results were shuffled across all test time points for each training time point, after which the same analysis as specified above was conducted to obtain new beta values for each quadrilateral. This process was repeated 100000 times to have a distribution of summed beta for each quadrilateral, from which the 95% thresholds were derived. The quadrilateral that had the summed beta values above the threshold and at the same time had the largest values among all quadrilaterals would be regarded as the one that significantly characterizes the representational generalization profiles.

## Results

### Experimental procedure and time-resolved RSA-based decoding analyses

Thirty-one human participants performed an auditory delayed-match-to-sample WM task while their 64-channel EEG activities were recorded. In each trial, during the encoding period, participants were presented with a 3-tone auditory sequence (Figure 1A) to memorize. During the maintaining period, a cuing tone that shared the same pitch as one of the three tones in the sequence was presented, and participants needed to maintain its corresponding ordinal position (1^st^, 2^nd^, 3^rd^) and made key pressing report during the subsequent retrieval period. Crucially, a neutral impulse, i.e., a white circle containing a symbolic ‘0’, was shown on the screen following the cuing tone, aiming to transiently perturb the WM system so that the retained ordinal position (1^st^, 2^nd^, 3^rd^) could potentially be reactivated. As shown in Figure 1A (right), participants performed relatively well in the task (86.88% ± 1.37%).

A time-resolved RSA-based decoding analysis (Fan et al., 2021; Kriegeskorte et al., 2008) was conducted on EEG activities to evaluate the neural representation of tone frequency and ordinal position, respectively, throughout the encoding and maintain phases (see details in Materials and Methods). The rationale of the RSA analysis is that the neural representational dissimilarity is proportional to the factor-of-interest dissimilarity, based on which the regression between the design dissimilarity matrix and the neural dissimilarity matrix was calculated to represent the decoding strength (beta).

### Concurrent neural representations of frequency and ordinal position during WM encoding

During the encoding period, each tone in the 3-tone sequence could be characterized by two factors, i.e., tone frequency (f1, f2, f3, f4, f5) and ordinal position (1^st^, 2^nd^, or 3^rd^). Figure 1B (middle panel) plots the time-resolved neural representational dissimilarity as a function of physical dissimilarity (y axis) after the onset of each tone, for tone frequency (left) and ordinal position (right). It is clear that the neural dissimilarity is minimum at center when the physical dissimilarity is also minimum, and increases for greater physical dissimilarity, supporting that the neural response indeed contained information for both tone frequency and ordinal position. Figure 1B (lower panel) shows the time-resolved regression coefficients (beta) for the two factors, further confirming that each tone in the sequence carries two neural codes in parallel, one for its pitch frequency (content; Figure 1B, left) and one for its ordinal position in the sequence (structure; Figure 1B, right) during the encoding period.

### Reactivation of ordinal position information during WM maintenance

We next probed memory information during maintenance when the 3-tone sequence is ‘silently’ retained in WM (Note that the decoding performance dropped to chance level after each auditory tone; Figure 1B). Specifically, the maintaining period was designed to contain two triggering events (Figure 2, upper, dotted box), a cuing tone indicating which ordinal position would be retrieved later and a task-irrelevant neutral visual impulse that is designed to potentially reactivate ordinal position information during retention.

**Figure 2.**
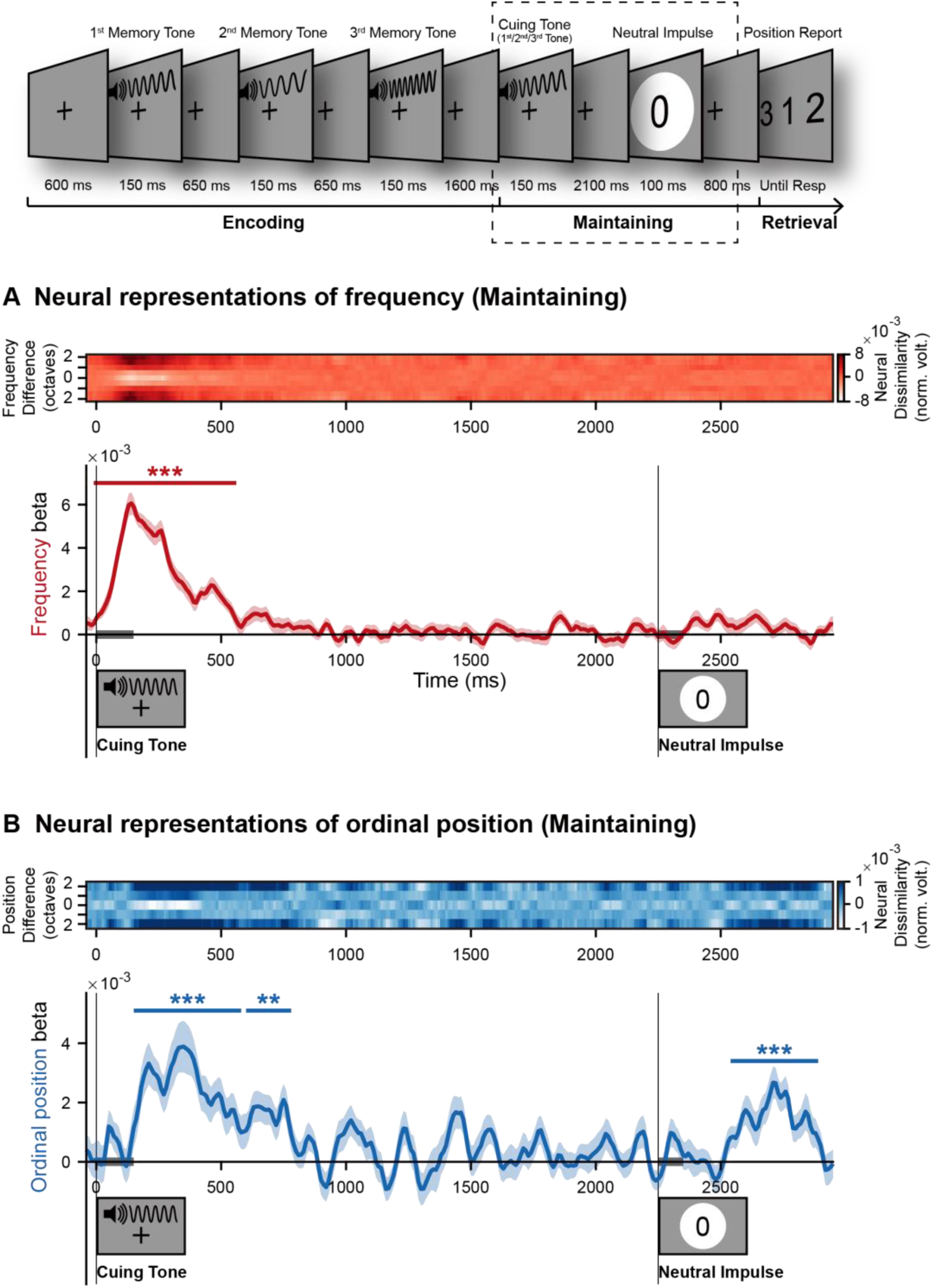
Memory reactivation during retention. Upper: dotted box indicates the maintaining period for decoding analysis. (A) Upper: grand average (N = 31) time-resolved neural representational dissimilarity (mean-centered) as a function of physical dissimilarity (y-axis) over time (x-axis) after onset of cuing tone, for auditory frequency. Lower: grand average (mean ± SEM) time-resolved decoding performance (regression between neural representational dissimilarity and physical dissimilarity) for tone frequency over time. Gray horizontal bar indicates triggering event presentation (cuing tone and neutral impulse). (B) The same as (A) but for ordinal position. (Shaded area represents ± 1 SEM across participants. ***: p < 0.001; **: p < 0.05; corrected using cluster-based permutation test, cluster-forming threshold p < 0.05).

Figure 2A plots the time-resolved neural representational dissimilarity (upper) and decoding performance (lower) for content (tone pitch) throughout retention. First, the tone frequency information could be successfully decoded from neural response to the cueing tone (Figure 2A), which is fully expected as the cuing tone by itself carries the pitch. Meanwhile, the following neutral impulse failed to elicit any pitch-related reactivations (Figure 2A). Since the main goal of the current experiment is to develop efficient triggers to reactivate ordinal position information, we will focus on the ordinal position reactivation in the following analysis.

Interestingly, both the cuing tone and neutral impulse successfully triggered the ordinal position representations (Figure 2B). Notably, the two triggering events – both the cuing tone and the neutral visual impulse – did not carry any explicit ordinal position information (1^st^, 2^nd^, or 3^rd^) in the stimulus. For instance, a cuing tone, although varied in pitch frequency in each trial, is not accompanied by any overt position tag when presented, and the following neutral visual impulse kept the same across all trials. Thus, the emergence of ordinal position representation after the two triggering events could not arise from any stimulus sensory characteristics (see Discussion for other interpretations)

Essentially, the ordinal position, once reactivated from WM system, was not sustained throughout retention but instead entered the ‘activity-silent’ state soon. Specifically, as shown in Figure 2B, the position representation that was reactivated by the cuing tone gradually decayed to baseline level after approximately 800 ms, and reemerged after the neutral visual impulse. In sum, ordinal position information ‘silently’ stored in WM could be reactivated intermittently throughout WM retention whenever efficient triggers are presented.

### Ordinal position reactivation during retention correlates with WM behavior

Having established the neural representations of ordinal position during encoding and maintaining periods, we next examined their WM behavioral correlates. Specifically, we divided all the participants into high- and low-performance groups (15 participants in each group) based on WM performance accuracies, and then compared the structure (i.e., ordinal position) neural representations during encoding and retention, respectively, between the two groups. As shown in Figure 3A, the high- and low-performance groups displayed similar ordinal position representation during encoding (Mann-Whitney U test, p = 0.500), suggesting that their varied memory performance was not due to encoding difference. In contrast, significant difference between the groups appeared when comparing the ordinal position reactivations after the cuing tone (Mann-Whitney U test; p < 0.001; Figure 2B) as well as after the neutral impulse (Mann-Whitney U test, p = 0.041; Figure 2C). Specifically, the high-performance group showed overall stronger reactivations compared to the low-performance group, confirming its genuine indexing of sequence structure storage in WM.

**Figure 3.**
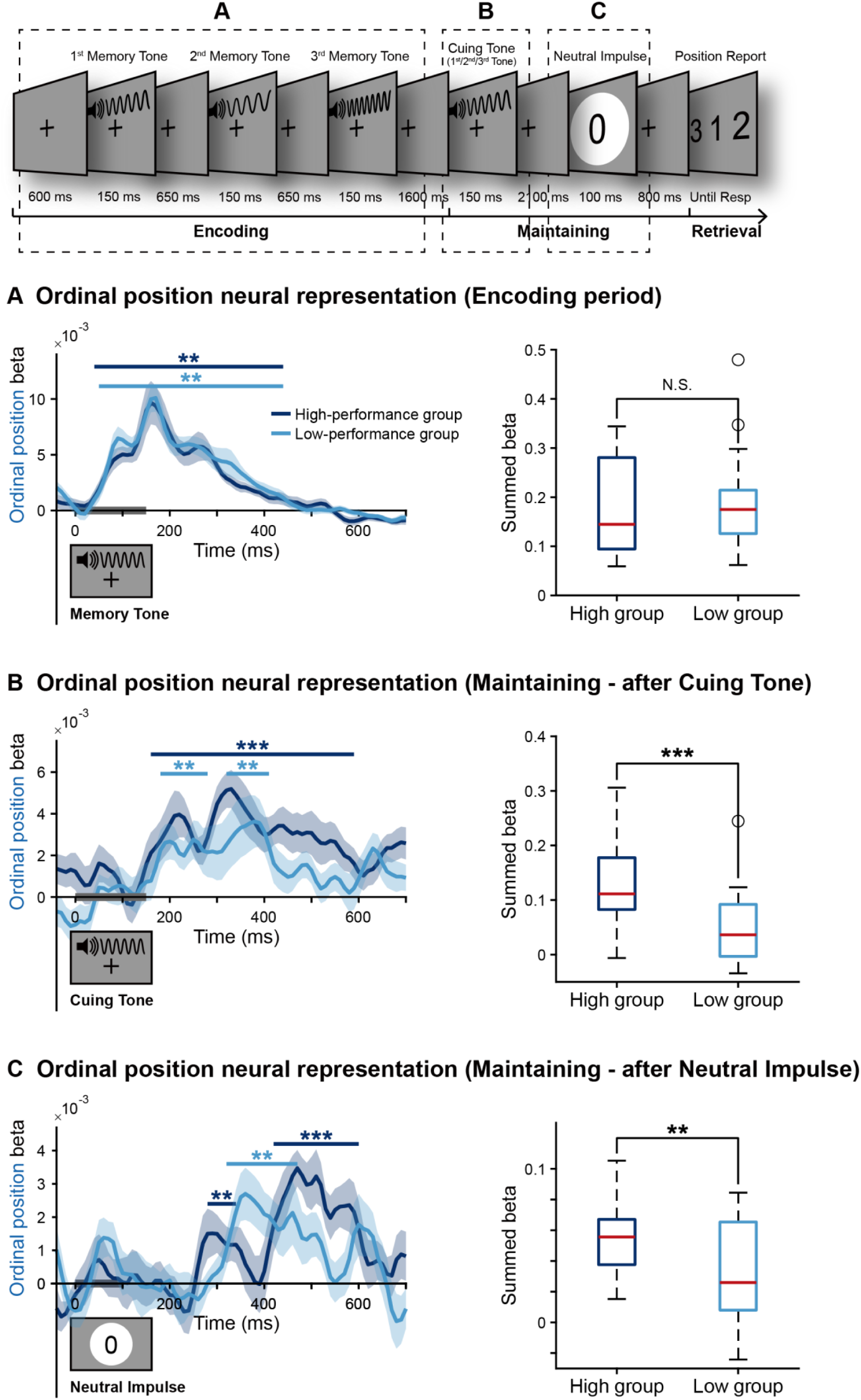
Behavioral correlates of ordinal position neural representations. Upper: dotted boxes indicate the epochs for decoding comparisons between high- and low-performance groups, corresponding to encoding (A), maintaining after cuing tone (B), and maintaining after neutral impulse (C). (A) Left: grand average (mean ± SEM) time-resolved ordinal position decoding performance during encoding period, for high-group (N = 15; dark blue) and low-performance groups (N = 15; light blue). Gray horizontal bar indicates memory tone presentation. Right: summed beta values that reached statistical significance throughout encoding period, for high-performance (dark blue) and low-performance groups (light blue). (B) Same as A but during maintenance after cuing tone. Gray horizontal bar indicates cuing tone presentation. (C) Same as A but during maintenance after neutral impulse. Gray horizontal bar indicates neutral impulse presentation. (All left panels: shaded area represents ± 1 SEM across participants. ***: p < 0.001; **: p < 0.05; corrected using cluster-based permutation test, cluster-forming threshold p < 0.05. All right panels: central red line indicates the median; bottom and top edges indicate the 25^th^ and 75^th^ percentiles, respectively. Outliers are indicated by hollow circle. Mann-Whitney U test; ***: p < 0.001; **: p < 0.05).

**Figure 4.**
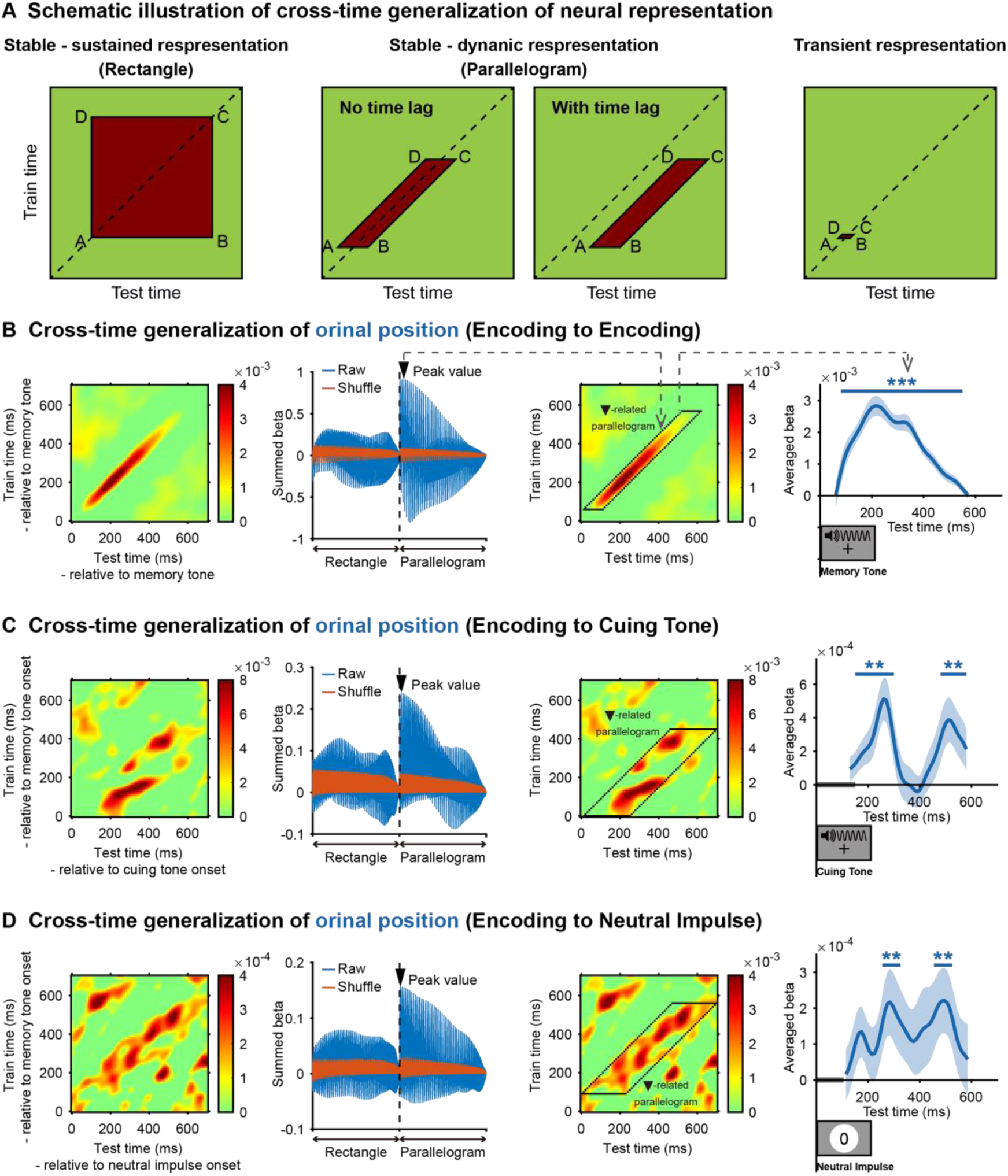
Cross-time representational generalization and “stable-dynamic” ordinal position code. (A) Schematic illustration of cross-time encoding-to-maintaining representational generalization that could account for stable and transient representation. Using encoding as training data (y-axis) and maintaining as testing data (x-axis). Left: “Stable-sustained” representation: information during encoding could be generalized to that during maintaining in a sustained way, displaying a horizontally-oriented rectangle in generalization results. Middle: “Stable-dynamic” representation: neural dynamic trajectory is repeated when reactivated during retention, without or with time delay, displaying a diagonally-oriented parallelogram in generalization results. Right: Transient representation: short burst of representation during retention. (B) Cross-time encoding-to-encoding representational generalization for ordinal position. Left: raw results. Middle left: characterization performance (summed beta values) for all horizontally-oriented rectangles and diagonally-oriented parallelograms (blue). Black triangle marks the quadrilateral with peak characterization performance above the permutation results (Orange). Middle right: The peak quadrilateral overlaid on the raw generalization results (black line). Right: the Beta time course extracted from the peak quadrilateral. (C) Same as B but for encoding-to-maintaining generalization after cuing tone. (D) Same as B but for encoding-to-maintaining generalization after neutral impulse. (All right panels in B, C, and D: Shaded area represents ± 1 SEM across participants. ***: p < 0.001; **: p < 0.05; corrected using cluster-based permutation test, cluster-forming threshold p < 0.05.)

### “Stable-dynamic” characteristics of ordinal position representation

We finally asked whether the neural code for ordinal position is the same or different between encoding and retention stages, by employing a cross-time encoding-to-maintaining generalization analysis (King & Dehaene, 2014). Although previous studies convergingly support stable representations for structure (Al Roumi et al., 2021; Fan et al., 2021; Kalm et al., 2017), the stable codes might arise from different possibilities, as illustrated in Figure 4A. For example, it might be specific temporal segment, in short or long duration, during encoding (y-axis, vertical span from A to D) that carries the structure code, which would be reactivated during retention in a sustained way (x-axis, horizontal span from A to B), referred to as “Stable-sustained representation” (Figure 4A, left). Alternatively, the code itself might be largely dynamic throughout the encoding period and follows a trajectory along the diagonal (see example in Figure 4B, right panel). Importantly, this dynamic pattern would repeat itself when reactivated during the delay period, without (Figure 4A, 2^nd^ from left; no time lag) or with time delay (Figure 4A, 2^nd^ from right; with time lag), referred to as “Stable-dynamic representation”. Finally, it is also possible that the ordinal position code and its reactivation occurs within a very short temporal bin (right panel, Figure 4A), namely “Transient representation”.

We first examined the temporal properties of ordinal position representation during encoding by performing a cross-time encoding-to-encoding generalization analysis. As shown in Figure 4B (left), the neural representation displays a diagonal coding profile, suggesting its highly dynamic characteristics, i.e., undergoing a highly dynamic transformation over time. Next, we asked whether the same dynamic trajectory would be repeated when reactivated during retention, and if so, we would expect a diagonally-oriented parallelogram profile in encoding-to-maintaining generalization results (Figure 4A, middle). Alternatively, if only specific segments of structure code during encoding is reactivated in a sustained manner during retention, we would expect a somewhat horizontally-oriented rectangle profile (Figure 4A, left). As shown in the left panels of Figure 4CD, the results indeed display a profile parallel to the diagonal, supporting the ‘stable-dynamic’ hypothesis. In other words, the time-resolved neural representation of ordinal position during encoding could be successfully generalized to that during retention over time, after both the cuing tone (Figure 4C) and the neutral impulse (Figure 4D).

Finally, we developed a new test to assess the statistical significance of the “stable-sustained” or “stable-dynamic” profile. Specifically, we established all possible horizontally-oriented rectangles (“stable-sustained”) and diagonally-oriented parallelograms (“stable-dynamic”) to cover the whole generalization matrix, and then calculated the summed beta values enclosed by each quadrilateral to represent its characterization performance (Figure 4BCD, middle left). A permutation test was then performed from which the quadrilateral that had the best and significant characterization performance was selected and then overlaid on the generalization matrix (Figure 4BCD, middle right). The generalization time course within the best quadrilateral was further extracted and plotted (Figure 4BCD, right). It turned out that the encoding-to-maintaining generalization results could best be captured via “Stable-dynamic” pattern (Figure 4CD)

Taken together, the ordinal position displays a ‘stable-dynamic’ representational format, that is, undergoing a dynamic neural trajectory over time, which reappears whenever reactivated during retention.

## Discussion

The goal of the present study is to examine the WM mechanism for the ordinal position of auditory sequences in human brains. To this end, we have developed new triggering events – cuing tone and neutral impulse – to successfully reactive the ordinal position information during WM retention. The behavioral relevance of the neural reactivation further confirms its genuine indexing of WM storage. Moreover, the ordinal position code is endowed with an interesting ‘stable-dynamic’ representational characteristic, which might facilitate efficient memory formation by embedding varied contents into stable-coded structures.

Memorizing a list of events in their occurring order is crucial in many cognitive functions (Averbeck et al., 2002; Dehaene et al., 2015; Hampton & Schwartz, 2004; Lashley, 1951; Pulvermüller, 2002; Tulving, 1972), and involves numerous brain regions, such as prefrontal cortex, parietal cortex, medial temporal lobe, and hippocampus (Amiez & Petrides, 2007; Foudil et al., 2020; Guidali et al., 2019; Lehn et al., 2009). To successfully retain a sequence in WM, two types of information – contents and the corresponding ordinal positions – need to be encoded and stored. Interestingly, ordinal structure and contents tend to show distinct storage characteristics, i.e., retained in different brain regions (Attout et al., 2019; Naya et al., 2017; Ninokura et al., 2004) and carried via different frequency bands of neural oscillations (Heusser et al., 2016; Hsieh et al., 2011; Kikumoto & Mayr, 2018; Yang et al., 2021). Sequence structure has been found to modulate how contents are represented in WM (Bellmund et al., 2021; Davachi & DuBrow, 2015; Huang et al., 2021). Here we extended previous results and provided new time-resolved neural evidence advocating explicit representations of ordinal position throughout the auditory WM process, i.e., during both encoding and retention periods.

It might be argued that participants simply retained verbal labels to denote the ordinal position and therefore the neural representation of ordinal position just reflects another type of content reactivation, akin to the pitch reactivation shown previously (Fan et al., 2021). Our results indeed do not support the interpretation, given that the neural characteristics of ordinal position are quite different from that of contents. Specifically, the ordinal position code displays a stable neural trajectory when reactivated during retention, while previous studies show that content representation generally undergoes a dynamic transformation from encoding to maintenance (Rademaker et al., 2019; Yu et al., 2020). Moreover, neural coding of ordinal position has also been shown in animals that lack language abstraction abilities (Bakhurin et al., 2017; Ninokura et al., 2003; Terada et al., 2017; L. Wang et al., 2015), supporting its species-general characteristic beyond language labeling.

How is information retained in WM? ‘Activity-based’ view supports persistent neuronal firing during the delay period (Constantinidis et al., 2018; Fuster & Alexander, 1971; X.-J. Wang, 2001), while the ‘Activity-silent’ view posits that memory could be retained in synaptic weights of the WM network (Barak & Tsodyks, 2014; Mongillo et al., 2008). Our results, consistent with previous EEG studies (Huang et al., 2021; Wolff et al., 2020), show that WM representation is not sustained throughout retention, but displays transient reactivations and would re-enter the silent state shortly. Nevertheless, it is noteworthy the lack of sustained representation might be due to the low signal-to-noise ratio in noninvasive electrophysiological recordings, and our results thus could not exclude the possibility that intracranial recordings or more sophisticated decoding analysis might reveal sustained profiles during WM retention (Barbosa et al., 2021; Kornblith et al., 2017). In fact, a recent study showed that information, although not decodable from raw EEG signals, was present in the alpha-band neural oscillations (Barbosa et al., 2021). Thus, our studies could not distinguish active- or silent-state WM views; Instead, the main goal here is to develop efficient approaches to reliably access ordinal position representations during the delay period. The ordinal position reactivation might arise from transient perturbation of the activity-silent state, but also could result from amplification of sustained memory traces (Barbosa et al., 2021).

To access memory representations during the delay period, one approach is bringing information to the focus of attention of WM (Harrison et al., 2009; LaRocque et al., 2013; Lewis-Peacock et al., 2012; Sprague et al., 2016), and the other is using a neural impulse to transiently reactive WM information (Fan et al., 2021; Wolff et al., 2015), corresponding to top-down and bottom-up approaches, respectively. Both methods aim to access or re-awaken the retained information to more active states so that they could be efficiently detected. Here, we combined the two operations, using a cuing tone to index a particular ordinal position (top-down) and a neural visual impulse (bottom-up) to reactivate the stored information. Our results demonstrate that both the cuing tone and the neutral visual impulse could trigger ordinal position, and these reactivations are further related to WM behavior. Rather than examining the structure information during the encoding period when stimuli are physically presented (Kalm & Norris, 2014; Marshuetz, 2005; Summerfield et al., 2020), here we, for the first time, developed triggering events with neutral characteristics to access the ordinal position code during retention.

By examining the time-resolved representational courses, we demonstrate that the neural code for the ordinal position is not stationary but follows a dynamic neural trajectory. Why does the ordinal position information undergo a specific neutral trajectory over time? It is posited to reflect ongoing reciprocal interactions between the active state and synaptic-based hidden state after external inputs enter the system (Stokes, 2015). The initial state and the synaptic weights of a network that reflect memory storage would together determine the dynamic trajectory unfolded over time for specific input (Barbosa et al., 2020). Here the ordinal position codes display dynamic trajectory throughout the whole memory process, i.e., during encoding, after cuing tone, and after neutral impulse, also consistent with previous findings (Miller et al., 2018; Murray et al., 2017; Spaak et al., 2017). Indeed, dynamic coding might be more energetically economical, compared to persistent firing (Crowe et al., 2010; Meyers, 2018).

Finally, the ordinal position code displays intriguing ‘stable-dynamic’ characteristics, constituting a new format for the stable nature of sequence structure in WM (D’Argembeau et al., 2015; Fan et al., 2021; Kalm et al., 2017). In contrast, the content code fails to show the encoding-to-maintaining representational generalization (Barak et al., 2010; Lundqvist et al., 2016; Quentin et al., 2019; Trübutschek et al., 2017; Yue et al., 2019). Content and structure are postulated to be signified in a factorized manner (Al Roumi et al., 2021; Liu et al., 2019; Ninokura et al., 2004), thereby a sequence is stored as a combination of stable structure (e.g., ordinal position) and varied content (e.g., tone pitch), on a trial by trial basis. The stable characteristics of structure would potentially facilitate memory formation, as the structure keeps steady to incorporate varied contents (Behrens et al., 2018; Friston & Buzsáki, 2016; Shahnazian et al., 2021). Moreover, the stable nature of the ordinal position code might reflect a general principle for structure at a broad level, e.g., cognitive map (Aronov et al., 2017; Garvert et al., 2017; Knudsen & Wallis, 2021).

## Acknowledgments

This work was supported by the National Science and Technology Innovation 2030 Major Program 2021ZD0204103 to H.L., and National Natural Science Foundation of China (31930052) to H.L.

